# Bounds to Parapatric Speciation: A Dobzhansky-Muller incompatibility model involving autosomes, X chromosomes and mitochondria

**DOI:** 10.1101/104489

**Authors:** Ilse Höellinger, Joachim Hermisson

**Affiliations:** Mathematics and BioSciences Group, Faculty of Mathematics and Max F. Perutz Laboratories, University of Vienna, 1090 Vienna, Austria; Vienna Graduate School of Population Genetics, Vienna Austria

**Author notes:** **Postal adress:** Joachim Hermisson and Ilse Höllinger, Faculty of Mathematics, University of Vienna, Oskar-Morgenstern-Platz 1, 1090 Wien, Austria, Europe, **Telephone number:** +43/1/4277 50648 (J. Hermisson), +43/1/4277 50771 (I. Höllinger), **Email-adress:**.

**Keywords:** hybrid incompatibility, two-locus DMI, speciation-with-gene-flow, large X-effect, introgression on X and autosomes

## Abstract

We investigate the conditions for the origin and maintenance of postzygotic isolation barriers, so called (Bateson-)Dobzhansky-Muller incompatibilities or DMIs, among populations that are connected by gene flow. Specifically, we compare the relative stability of pairwise DMIs among autosomes, X chromosomes, and mitochondrial genes. In an analytical approach based on a continent-island framework, we determine how the maximum permissible migration rates depend on the genomic architecture of the DMI, on sex bias in migration rates, and on sex-dependence of allelic and epistatic effects, such as dosage compensation. Our results show that X-linkage of DMIs can enlarge the migration bounds relative to autosomal DMIs or autosome-mitochondrial DMIs, in particular in the presence of dosage compensation. The effect is further strengthened with male-biased migration. This mechanism might contribute to a higher density of DMIs on the X chromosome (large X-effect) that has been observed in several species clades. Furthermore, our results agree with empirical findings of higher introgression rates of autosomal compared to X-linked loci.

## 1 Introduction

Historically, speciation research has mostly focused on two idealized scenarios: allopatric speciation (complete geographic isolation of incipient species) and sympatric speciation (divergence of subpopulations in a common habitat) (Orr and Turelli, 2001; Coyne and Orr, 2004; Via and West, 2008). Both scenarios are simplifications of biological reality. While strict sympatry of incipient species seems to be an exception, there is abundant evidence for hybridization even among “good species” with viable and not completely sterile hybrid offspring (reviewed *e.g.* in Coyne and Orr, 2004; Mallet, 2008). Population genetic theory shows that even low levels of gene flow can strongly interfere with population differentiation (Felsenstein, 1981; Slatkin, 1987). This makes it inevitable to assess the impact of limited gene flow at various stages of the speciation process, a scenario commonly referred to as parapatric speciation.

The classical model for the evolution of postzygotic isolation barriers in allopatry is the (Bateson-)Dobzhansky-Muller model (DMM) (Bateson, 1909; Dobzhansky, 1936; Muller, 1942). The DMM assumes that new substitutions occur on different genetic backgrounds. When brought into secondary contact, these previously untested alleles might be mutually incompatible and form Dobzhansky-Muller incompatibilities (DMIs), thus reducing hybrid fitness and decreasing gene flow at linked sites. The emergence of species boundaries due to accumulation of DMIs in allopatry is well understood (Coyne and Orr, 1989; Orr and Turelli, 2001; Coyne and Orr, 2004). More recently, several studies have considered this process in parapatry (Gavrilets, 1997; Feder and Nosil, 2009; Agrawal et al., 2011; Bank et al., 2012; Wang, 2013; Lindtke and Buerkle, 2015). All support that the DMM provides a viable mechanism for the evolution of postzygotic isolation even in the presence of gene flow, although the bounds for maximum permissible migration rates can be quite stringent.

Empirically, there is widespread evidence for DMIs not only among recently diverged sister species (Maheshwari and Barbash, 2011; Presgraves, 2010; Sweigart and Flagel, 2014), but also segregating within species (Corbett-Detig et al., 2013). Hence, these authors argue that the genetic basis of reproductive isolation is continuously present within natural populations, rendering the independent allopatric evolution of newly incompatible substitutions obsolete.

While most theoretical studies focus on autosomal DMIs, empirical evidence points to a major role of sex chromosomes in speciation. Haldane’s rule (Haldane, 1922, reviewed in Coyne and Orr, 2004), states that in species with sex specific reduced hybrid fitness the affected sex is generally heterogametic. The *large X*-*effect* (Coyne and Orr, 1989, reviewed in Presgraves, 2008) expresses the disproportional density of X-linked incompatibility genes in postzygotic isolation. For example Masly and Presgraves (2007) report a higher density of incompatibilities causing hybrid male sterility on the X chromosome relative to autosomes in *Drosophila*. Equivalent findings exist of a *large Z-effect* in WZ-systems, such as birds, where WZ-females are heterogametic (Ellegren, 2009). Also cytoplasmic incompatibilities have been described (Ellison and Burton, 2008; Lee et al., 2008; Burton and Barreto, 2012; Barnard-Kubow et al., 2016).

A recent study by Bank et al. (2012) determined stability conditions and maximum permissible migration rates of autosomal two-locus DMIs in a continent-island framework. They distinguished two main mechanisms shaping the evolution of DMIs: selection against (maladapted) immigrants and selection against (unfit) hybrids, which lead to different dependence of maximum migration rates on the model parameters.

Prompted by the empirical observations described above, we extend the model by Bank et al. (2012) to general two-locus DMIs in diploids involving X chromosomes, autosomes, or mitochondria. We include sex-specific fitness effects, in particular, to account for the effect of dosage compensation of hemizygous X-linked genes in males. We also allow for sex-specific migration, as many species display differences in migration patterns for males and females Greenwood (1980).

Following Bank et al. (2012) we derive maximum migration bounds where DMIs can still originate in parapatry, or resist continental gene flow. In contrast to the autosomal case, we find that sex specific fitness- and sex-biased migration cause substantial differences in the maximum permissible rates and hence influence the prevalence of autosomal DMIs relative to X-linked and mitochondrial DMIs. Especially, we find that X-linkage of (nuclear or cytonuclear) DMIs together with dosage compensation and/or male-biased migration boosts migration bounds and thus enhances the evolution of X-linked DMIs, possibly contributing to a *large X*-*effect* and to reduced introgression probabilities of X-linked DMI loci.

## 2 Model and Methods

We consider a diploid, dioecious population with separate sexes (at 1:1 ratio) that is divided into two panmictic subpopulations, continent and island. (See Figure 1 and Figure B.1 in the Supporting Information (SI)). Both demes are sufficiently large that drift can be ignored (drift effects are discussed in SI Section A.3). They are connected by unidirectional sex-dependent migration at rate *m*^♀^ and *m*^♂^ from the continent to the island. We fix the average migration rate per individual, 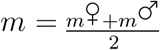, and define

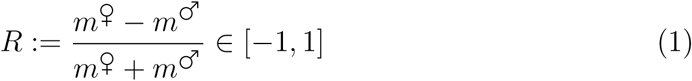

as a measure of sex-bias in migration. Sex-specific migration gives rise to distinct migration rates per allele for autosomes, X chromosomes, and mitochondria, *𝓂_𝒜_*, *𝓂_𝒳_*, *𝓂_𝒪_* (Eqs. (B.13)-(B.15)). For *−*1 ≤ *R* < 0 migration is male-biased and we obtain *𝓂_𝒜_* > *𝓂_𝒳_* > *𝓂_𝒪_*. In contrast, for 0 < *R* ≤ 1 migration is female-biased, resulting in *𝓂_𝒜_* < *𝓂_𝒳_* < *𝓂_𝒪_*.

**Figure 1:**
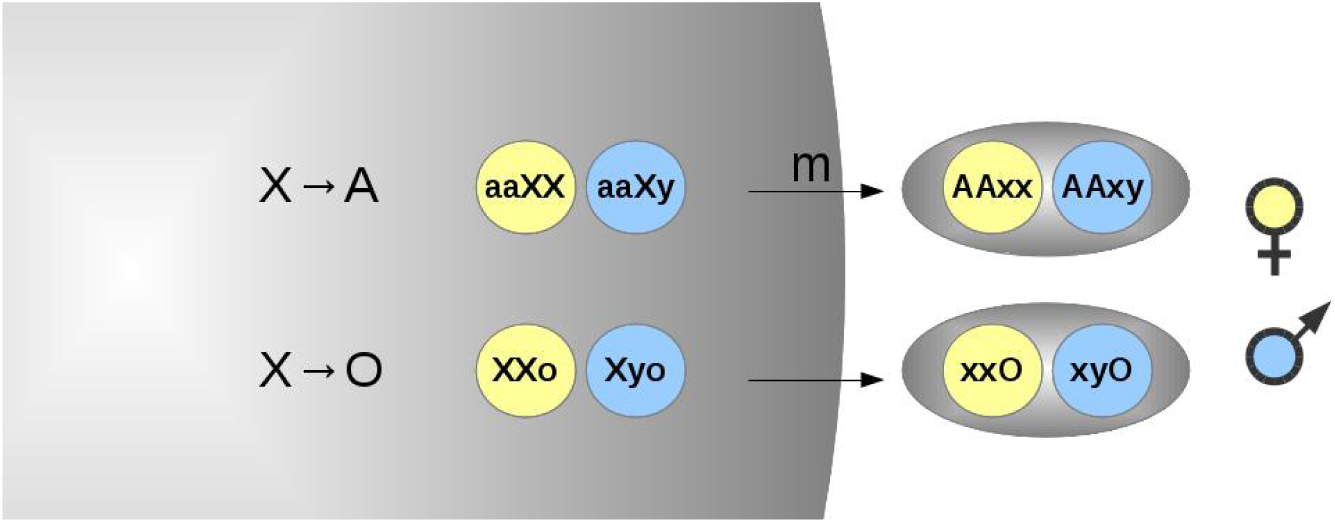
Schematic model. The population inhabits a continent (left) and an island (right), which are connected by unidirectional migration at rate *m*. The figure shows two out of eight genomic architectures investigated: an X-autosome DMI (upper line) and a cytoplasmic DMI between X and mitochondrion (lower line). Genotypes of female residents are depicted by yellow circles and males by blue circles, respectively. The capital letters denote incompatible alleles, which reduce hybrid fitness.

### The DMI

The incompatibility is formed by two unlinked biallelic loci, situated on autosomes *𝒜*, X chromosomes *𝒳*, or in the mitochondrial genome (cytoplasmic organelle) *𝒪*, (cf. Table 1). Both sexes are diploid for autosomes and haploid for the mitochondrial locus. Males are hemizygous for the X chromosomes, whereas females are diploid. The continent is monomorphic for the continental (geno-)type and only acts as source of migrants for the island. Our analysis focuses on the evolutionary dynamics on the island. A stable DMI corresponds to a stable equilibrium on the island where all four alleles are maintained (a two-locus polymorphism), including the pair of incompatible alleles (indicated by capital letters in Table 1).

**Table 1:** Each genomic architecture is defined by a continental (geno-)type (third column) and an island (geno-)type (fourth column). Mutually incompatible pairs of DMI-alleles are denoted by capital letters. We call the immigrating DMI-allele *continental allele* and its resident incompatible partner *island allele*. The name of each model in the first column is constituted by “the continental allele → the island allele”. The A→A-model corresponds to the model by Bank et al. (2012).

| Model | DMI | continental type ($\varphi, \sigma$) | island type ($\varphi, \sigma$) |
| --- | --- | --- | --- |
| $A \rightarrow A$ | $\mathcal{A}-\mathcal{A}$ | aaBB | AAbb |
| $X \rightarrow A$ | $\mathcal{A}-\mathcal{X}$ | aaXX, aaXy | AAxx, AAxy |
| $A \rightarrow X$ | $\mathcal{A}-\mathcal{X}$ | AAxx, AAxy | aaXX, aaXy |
| $X \rightarrow X$ | $\mathcal{X}-\mathcal{X}$ | $X_1X_1x_2x_2, X_1x_2y$ | $x_1x_1X_2X_2, x_1X_2y$ |
| $A \rightarrow O$ | $\mathcal{A}-\mathcal{O}$ | AAo | aaO |
| $O \rightarrow A$ | $\mathcal{A}-\mathcal{O}$ | aaO | AAo |
| $X \rightarrow O$ | $\mathcal{X}-\mathcal{O}$ | XXo, Xyo | xxO, xyO |
| $O \rightarrow X$ | $\mathcal{X}-\mathcal{O}$ | xxO, xyO | XXo, Xyo |

We model genotypic fitness as the sum of direct allelic fitness and epistasis. Hence any given allele contributes directly to genotype fitness, where is can be locally or globally adapted, and additionally via epistais if it is incompatible with other alleles in the same genotype. We set the (Malthusian) fitness of genotypes containing no incompatible alleles (only lower case letters) in both sexes to 0. For simplicity, we assume no dominance of the single-locus effects, but we allow for dominance or recessitivity of the incompatibility.

### Different genomic architectures of DMIs

We define the fitness of an arbitrary female genotype as

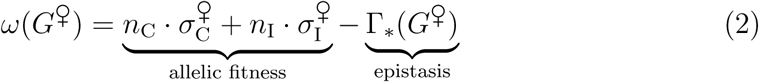

or for a male genotype as

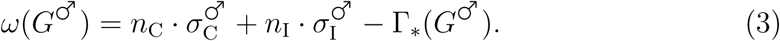

The allelic fitness is captured by the selection coefficient σ^♀^,σ^♂^ (for females and males) and weighted with the respective number of incompatible alleles, *n*_C,I_ ∈ {0, 1, 2} in a given genotype. To match the locus effects of haploid mitochondrial genes to autosomes, we set *n*_C,I_ ∈ {0, 2} for the absence or presence of the single incompatible allele in this case.

We assume σ^♀^ = σ^♂^ for autosomes and organelles, but for X-linked alleles the fitness effect may be enhanced in males, σ^♀^ = (1 + *D*)σ^♂^, where *D* ∈ {0, 1} measures dosage compensation (see below). The contribution of epistasis to hybrid genotype fitness can be summarized by an epistasis vector Γ_*_, for each model (_*_), detailed in Table 2

**Table 2:** The table shows all possible hybrid genotypes with DMIs (second column) and corresponding fitness cost, parametrized by the entries of the epistasis vector (third column). The strength of the incompatibility depends on the number of incompatible alleles in the genotype. Plausibly, the strength increases with the number of incompatible pairs, which can be 1, 2, or 4 (Turelli and Orr, 2000). We focus on two particular epistasis schemes, one with a codominant DMI 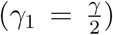 with fitness cost proportional to the number of incompatible pairs and one with a recessive DMI (γ_1_ = 0) where the fitness cost is zero if there is still a pair of compatible alleles in the genotype. The strength of X-linked incompatibilities in males depends on dosage compensation, captured by *D* ∈ {0, 1}.

| DMI | hybrids: $\varphi, \sigma$ | epistasis vector $\Gamma_*$ |
| --- | --- | --- |
| $\mathcal{A}\text{-}\mathcal{A}$ | $\varphi$: AaBb, AaBB, AABb, AABB<br>$\sigma$: AaBb, AaBB, AABb, AABB | $\Gamma_{\mathcal{A}\mathcal{A}} = (\underbrace{\gamma_1, \gamma, \gamma, 2\gamma}_{\varphi}, \underbrace{\gamma_1, \gamma, \gamma, 2\gamma}_{\sigma})$ |
| $\mathcal{A}\text{-}\mathcal{X}$ | $\varphi$: AaXx, AaXX, AAXx, AAXX,<br>$\sigma$: AaXy, AAXy | $\Gamma_{\mathcal{A}\mathcal{X}} = (\gamma_1, \gamma, \gamma, 2\gamma, (1 + D)\frac{\gamma}{2}, (1 + D)\gamma)$ |
| $\mathcal{X}\text{-}\mathcal{X}$ | $\varphi$: X<sub>1</sub>x<sub>1</sub>X<sub>2</sub>x<sub>2</sub>, X<sub>1</sub>x<sub>1</sub>X<sub>2</sub>X<sub>2</sub><br>X<sub>1</sub>X<sub>1</sub>X<sub>2</sub>x<sub>2</sub>, X<sub>1</sub>X<sub>1</sub>X<sub>2</sub>X<sub>2</sub>, $\sigma$: X<sub>1</sub>X<sub>2</sub>y | $\Gamma_{\mathcal{X}\mathcal{X}} = (\gamma_1, \gamma, \gamma, 2\gamma, (1 + 3D)\frac{\gamma}{2})$ |
| $\mathcal{A}\text{-}\mathcal{O}$ | $\varphi$: AaO, AAO, $\sigma$: AaO, AAO | $\Gamma_{\mathcal{A}\mathcal{O}} = (\gamma, 2\gamma, \gamma, 2\gamma)$ |
| $\mathcal{X}\text{-}\mathcal{O}$ | $\varphi$: XxO, XXO, $\sigma$: XyO | $\Gamma_{\mathcal{X}\mathcal{O}} = (\gamma, 2\gamma, (1 + D)\gamma)$ |

**Strength of the incompatibility: The epistasis vector**

### Dosage compensation

Dosage compensation can be related to different mechanisms. For example, in the model organism *Drosophila melanogaster* the expression of the X chromosome is doubled in males. An alternative mechanism has evolved in mammals, where one X chromosome is randomly inactivated in females (Payer and Lee, 2008). Finally, in birds dosage compensation seems to be incomplete, as some genes show elevated expression levels in homogametic ZZ-males compared to heterogametic females, whereas other genes are dosage compensated (Ellegren et al., 2007; Graves et al., 2007).

Our model allows for arbitrary sex-dependence of allelic and epistatic effects, but we focus on dosage compensation of the hemizygous X chromosome in males as a key biological mechanism. We model fitness for any X-linked allele in hemizygous males in two ways (Charlesworth et al., 1987):

- *No dosage compensation, D* = 0: A single copy of an X-linked allele has the same allelic (σ^♀^ = σ^♂^) and epistatic effects in hemizygous males as in females.
- *Full dosage compensation, D* = 1: Hemizygosity of the X chromosome is compensated in males, and a single X-linked allele has the same effect as a homozygous pair of X chromosomes in females (allelic selection coefficient: (σ^♀^ = 2σ^♂^).

With random deactivation of X in females we naturally obtain a codominant DMI in our model since (average) heterozygous fitness is equal to the mean of the homozygous fitnesses in this case.

### Dynamics of the general model

For our analytical treatment, we assume weak evolutionary forces, such that linkage equilibrium among both loci and Hardy-Weinberg-proportions can be assumed. It is then sufficient to track the frequencies of the continental allele *p*_C_ and the island allele *p*_I_ on the island. We test this approximation for stronger selection by numerical simulations in SI Section A.2.

For each genomic architecture (Table 1) we derive a pair of differential equations in continuous time (see **Box 1**). For the case of an X→A DMI, in particular, *p*_C_ measures the frequency of the incompatible X allele that immigrates from the continent and *p*_I_ the frequency of the incompatible autosomal allele on the island.

#### Box 1

Dynamics of the continental allele frequencies *p*_C_:

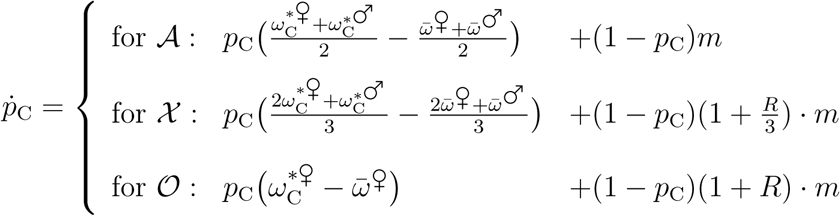

Dynamics of the island allele *p*_I_:

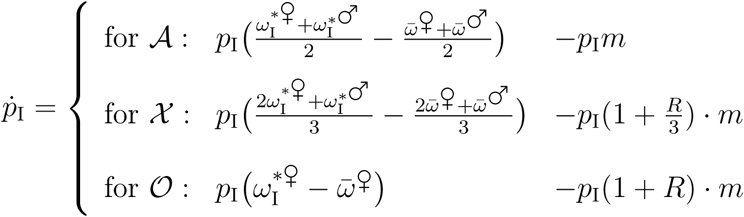

Marginal fitness 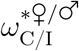 and mean fitness 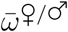 for each sex are functions of genotype fitness (consult SI Eqs.(B.9),(B.10) for explicit expressions). Sex-bias in migration *m* is measured by *R* (Eq.(1)).

We obtain:

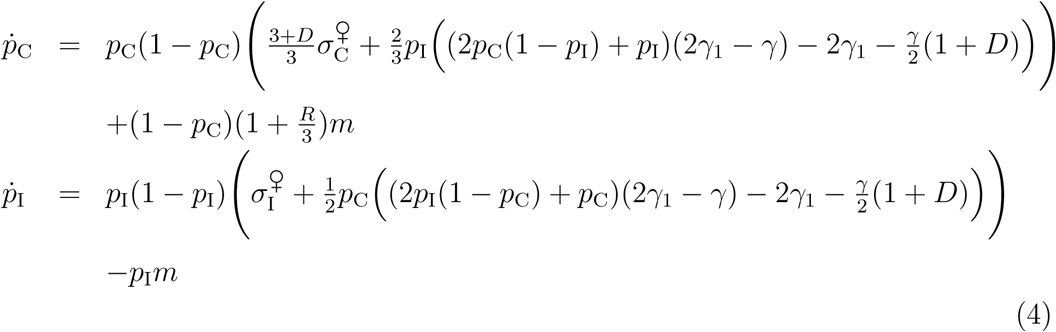

We see that with dosage compensation (*D* = 1), the X-linked allelic fitness is increased 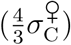, because a single X-allele in males now acts as strongly as two X-alleles in females. Similarly, dosage compensation increases the term due to epistasis in males 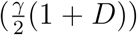 affecting both the X-linked allele and the autosomal allele. Sex-biased migration, quantified by *R* (see Eq. 1), affects only the X-linked allele, as males are hemizygous X carriers. Parameterizations for all other cases are provided in the SI Section B.1.2.

### The codominant model

If the effect of the incompatibility is additive, such that it is proportional to the number of incompatible pairs in a genotype (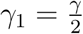 in Table 2), the model simplifies greatly. For the X→A model, in particular,

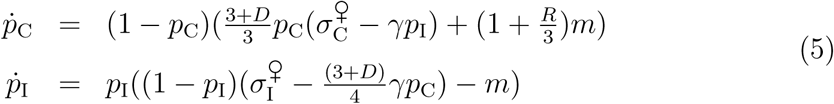

see SI Eqs. (B.34) for the other models.

### Evolutionary histories

A parapatric DMI can evolve via different routes, depending on the timing and geographic location of the origin of the two mutations. Following Bank et al. (2012), we distinguish five histories: For *secondary contact*, both substitutions occur during an allopatric phase and can originate in any order. In contrast, if the substitutions originate in the presence of gene flow, the timing matters and we obtain four further scenarios: for a *continent-island* DMI we have the first substitution originating on the continent and the second on the island. Analogously, there are *island-continent*, *island-island*, and *continent-continent* scenarios. Note that the first two scenarios lead to *derived-derived DMIs*, with one substitution in each deme, whereas the last two lead to *derived-ancestral DMIs*, where both substitutions occur in the same deme. In all cases we refer to the immigrating incompatible allele as the *continental allele* and to the resident, incompatible allele as the *island allele*. All five evolutionary histories lead to the same dynamics (as given in Box 1) upon appropriate relabeling of genotypes, where different histories correspond to different initial conditions (see SI Section B.2.5 and “Mapping of evolutionary histories” below).

## 3 Results

Our analytical analysis of the dynamical system in **Box 1** is presented in comprehensive form in SI B. It comprises on the following steps. For the general model (0 ≤ γ_1_ < γ), we determine all boundary equilibria and conditions for their stability. Instability of all boundary equilibria implies a protected polymorphism at both loci. Excluding cycling behavior, this is a sufficient condition for a globally stable DMI that will be reached from all starting conditions (all evolutionary histories). An internal stable equilibrium (DMI) can also coexist with a stable boundary equilibrium. In this case, the DMI is only locally stable and will only be reached from favorable starting conditions. Necessary and sufficient conditions for the existence of (locally or globally) stable DMIs can be derived for weak migration by means of perturbation analysis: A stable DMI results if the monomorphic boundary equilibrium (*p*_I_ = 1, *p*_C_ = 0) is stable for *m* = 0 and is dragged inside the state space for small *m* > 0. For codominant DMIs, also the internal equilibria can be assessed analytically and conditions for stable DMIs follow from a bifurcation analysis. For the recessive DMIs, we complement our analytical results by numerical work to derive stability conditions for locally stable DMIs under stronger migration.

Below, we summarize the key results for the general model. This is followed by a detailed analysis of the codominant model. In the supplement we added continuative results, first for the recessive model in SI Section A.1. Second, SI Section A.2 contains simulation results to assess the effects of linkage disequilibrium (LD), which is relevant for very strong incompatibilities. Third, we present simulations for finite populations and analyze how migration limits are affected by genetic drift in SI Section A.3. Finally, in SI Section A.4 we calculate adaptive substitution rates for autosomes and X chromosomes with gene flow and derive conditions on dominance favoring the *faster X-effect*, described by Charlesworth et al. (1987).

### Stable equilibria: global and local stability of DMIs

The model has three boundary equilibria: A monomorphic state, where the continental genotype swamps the island, which is always reached for strong migration. Furthermore, two single locus polymorphisms (SLPs) where one locus is swamped, but the other is maintained polymorphic. There is at most one stable internal equilibrium, corresponding to a DMI. It can either be globally or locally stable. In the latter case, one of the boundary equilibria is also locally stable and it depends on the evolutionary history which equilibrium is reached. We therefore obtain two migration thresholds 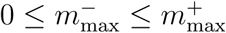:

- For migration rates 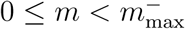, a globally stable DMI, that is reached for all evolutionary histories.
- For migration rates 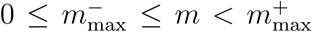, the dynamics are bistable and yield a locally stable DMI. Hence, only certain evolutionary histories permit its evolution, but any existing DMI will be maintained.
- For migration rate 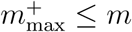 no stable DMI exists.

### Mapping of evolutionary histories

Every evolutionary history maps to a distinct initial condition (SI Section B.2.5 for results and proofs). As in Bank et al. (2012), we find three permissive histories that always result in the evolution of a stable DMI for 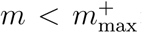: *secondary contact*, *island-continent*, and *continent-continent*. In all these cases, the second substitution occurs in a deme where the incompatible first substitution is not (yet) present. In contrast, the evolution of a stable DMI in parapatry is more difficult for an *island-island* or *continent-island* substitution history. Here, the second substitution needs to invade on the island despite of competition of the incompatible allele. We need 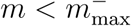 for a DMI to originate under these circumstances.

### Necessary conditions for the existence of a stable DMI

Based on previous results for the model without migration (Rutschman, 1994) or without epistasis (Bürger and Akerman, 2011), and in accordance to Bank et al. (2012), we find that with epistasis and increasing migration a stable DMI can only exist if the island allele is beneficial and its sex-averaged selection coefficient exceeds migration. Furthermore, any averaged selective advantage of the continental allele must be outweighed by averaged epistasis. For example for X→A, we obtain 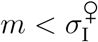 and 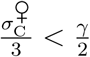. Consult Eqs. B.29,B.31 and Table B.3 for different terms for each model and the SI Sections B.1.3, B.1.6 for proofs.

### 3.1 Nuclear codominant DMIs

We obtain full analytical solutions for the maximum migration bounds 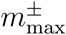 (B.2.4). Below, we discuss how these rates depend on the various genetic architectures, sex dependence of fitness and migration, and on dosage compensation. Figures 2 and 3 compare the 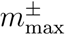 for the different types incompatibilities among nuclear genes: DMIs among autosomal genes (A→A), DMIs among X and autosomes, with either the incompatible X allele immigrating from the continent (X→A) or the autosomal locus (A→X), and DMIs among two X-linked loci (X→X). Figure 2 assumes full dosage compensation of X-linked alleles in males, Figure 3 treats the case without dosage compensation.

**Figure 2:**
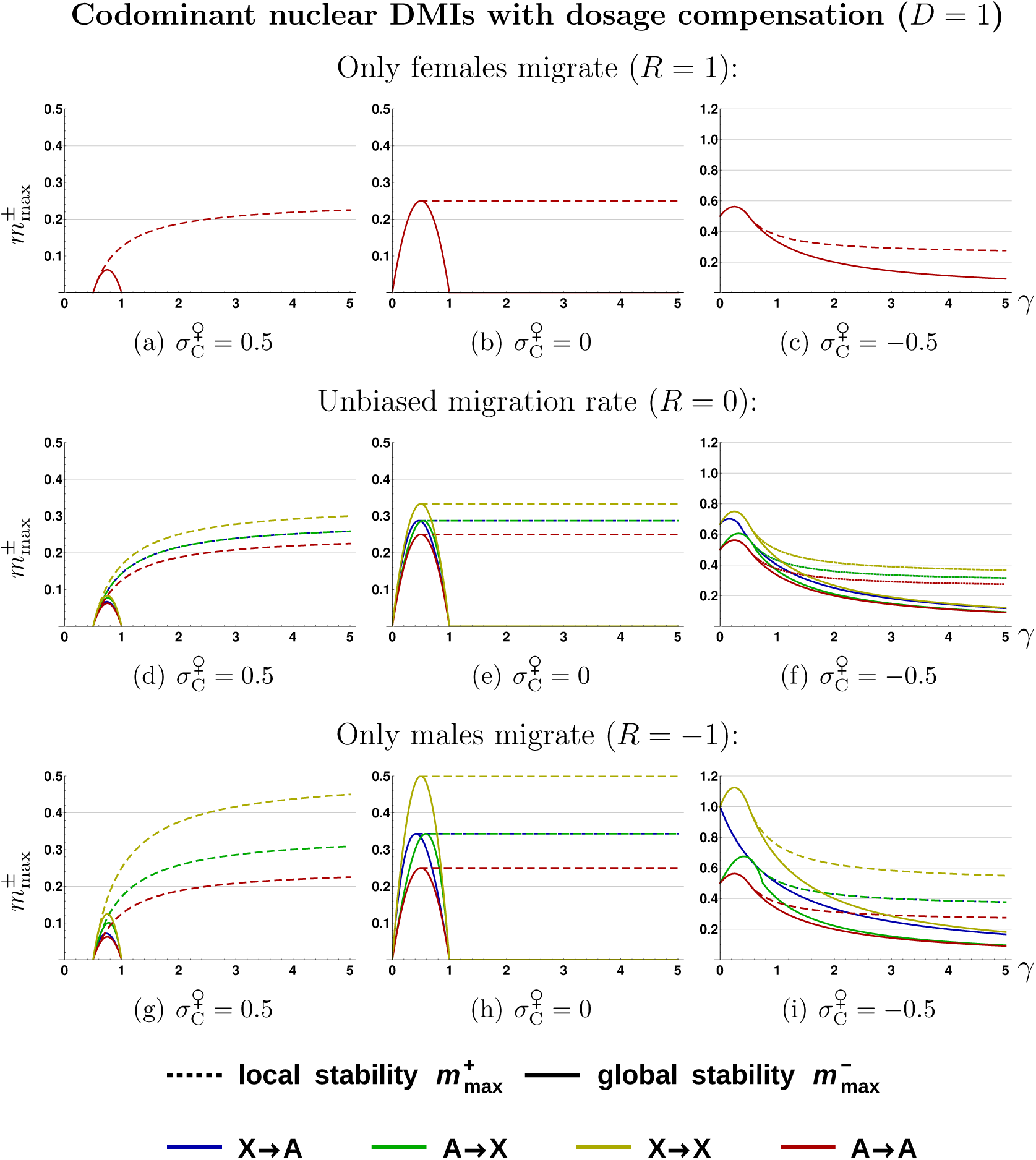
Codominant nuclear DMIs with dosage compensation, *D* = 1. The columns show 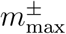 as a function of the strength of epistasis γ for beneficial 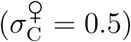, neutral 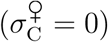, and deleterious 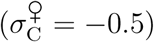 effect of the immigrating allele. All quantities in the figure 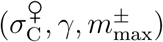 are measured relative to the fitness effect of the island allele, which is normalized to 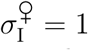. Note the different scaling of the y-axis in the third column. Strong differences between 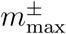 in the various models occur if migration rates are sex-biased. For female-biased migration 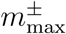 coincide for all four models. With increasing proportion of male migrants (top to bottom), migration pressure on the X chromosome is reduced and differences among the models appear. All bounds 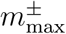 are derived analytically, see Eqs.(B.40),(B.42).

**Figure 3:**
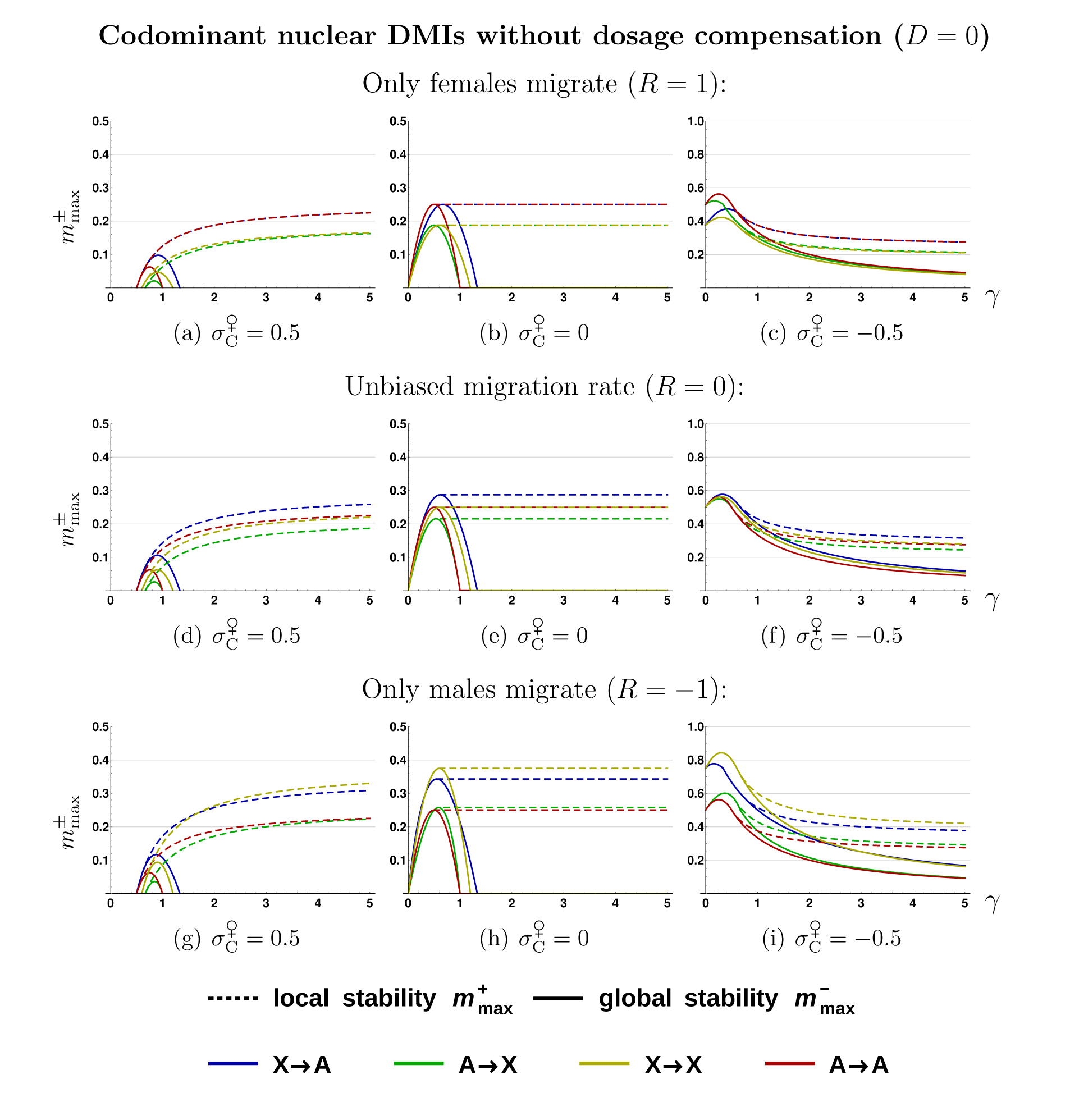
Codominant nuclear DMIs without dosage compensation, *D* = 0. Without dosage compensation the ploidy differences between the autosomes and the X chromosome are unmasked, inducing strong asymmetry between the *𝒜*-*𝒳*-models. this leads to a larger effect per allele. All bounds 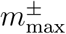 are derived analytically, see Eqs.(B.40),(B.42). See also Figure 2 for further explanations. Note the different scaling of the y-axis in the third column.

#### Selection against hybrids and against immigrants

Following Bank et al. (2012), we can distinguish two main selective forces maintaining a DMI in the face of gene flow. If the continental allele is beneficial on the island (first column of Fig. 2 and 3), a polymorphism at the respective locus can only be maintained by hybrid formation and selection against the immigrating allele is due to hybrid inferiority (“selection against hybrids”). This type of selection will only be effective as long as the immigrating allele is rare. Once the migration pressure is so high that the immigrating continental allele is in a majority, incompatibility selection rather works against the resident allele on the island. Consequently, we expect a large bistable regime with 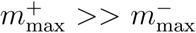 and a small region with global stability, as can indeed be seen for all types of DMIs with a beneficial continental allele. Note also that 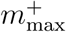 increases with γ, as should be expected if hybrid incompatibility, *i.e.* epistasis, is the sole cause of (local) stability.

In contrast, with a deleterious immigrating allele (third column of Fig. 2 and 3), a DMI can also be maintained by “selection against immigrants” for small values of epistasis, or via a combination of the two selective forces (selection against hybrids and immigrants) with stronger epistasis. If selection against immigrants predominates, maintenance of the DMI is driven by local adaptation. The fitness advantage of the resident allele depends on its direct effect and the dynamics will usually be frequency independent. Therefore we obtain no or only a small bistable regime, with 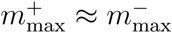. For stronger epistasis, selection against hybrids becomes more important, leading to a relative increase of the bistable regime. The main effect of epistasis now is that it promotes swamping of the island allele: 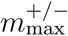 decreases with epistasis. In the case of a neutral immigrating allele, the observed migration bounds exhibit an intermediate pattern.

#### Sex-biased migration

To understand the differences among the DMI architectures, we take the case of full dosage compensation and strict female-biased migration (*R* = 1) as a starting point (Fig. 2(a)-(c)). In this case, all curves for 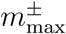 for the different models collapse onto a single one. Indeed, if only females migrate, the number of migrating X chromosomes and autosomes is equal. Full dosage compensation balances any direct and epistatic effects of loci with different ploidy levels. Consequently, the corresponding Eqs. (B.34) differ only by a constant factor.

If also males migrate (Fig. 2(d)-(i)) genomic architectures involving an X chromosomes experience effectively lower migration rates of the X and hence increasing 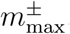. Male-biased migration boosts 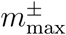 most effectively for X→X, as both loci experience reduced migration pressure. For unbiased migration, 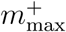 of X→X relative to A→A DMIs increases by 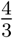 (the autosome-X ratio), and doubles for pure male migration (corresponding to the 1:2 X-autosome ratio among migrants in this case).

The migration bounds 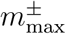 for the A→X and X→A DMIs are intermediate between the A→A and X→X DMIs. Our analytical results (see B.2) show that the upper limit of the bistable regime (*i.e.*, the value of 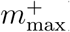) is identical for the A→X and the X→A models with dosage compensation. However, the limits for global stability, 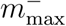, can differ, which can be understood as follows:

For pure selection against migrants (no epistasis γ → 0, and σ_C_ < 0, right column in Fig. 2), increased male migration reduces the effective migration pressure on the X chromosome. This leads to a corresponding increase in the migration bound 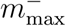 (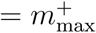 in this case) for all DMIs that are lost for 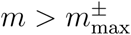 because of swamping at an X-locus. This is clearly always the case for X→X DMIs, but also for the X→A model, as long as |σ_C_| < |σ_I_| (as in our example: X fixes before A is lost). In contrast, for the A→A and the A→X model (if |σ_C_| < |σ_I_|) the DMI is lost due to swamping at the autosomal locus (continental locus in these cases). Increased male bias in migration therefore does not change the migration bound 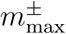 in these cases.

For strong epistasis (with a deleterious immigrating allele), where the direct locus effects are less important, it is always the incompatible island allele that cannot invade on the island for migration rates 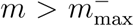. Here, any incompatible island allele that interacts with an X allele has an advantage from male-biased migration since it feels less gene flow from the competing X. This can be seen for the 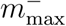 lines in Figure 2(f),(i): While the migration bound is increased for the X→A model (and the X→X model) over the whole range of epistasis, it converges to the value of the autosomal DMI for the A→X model.

#### No dosage compensation

In Figure 3 we investigate migration bounds without dosage compensation, such that the differences in ploidy between autosomes and X chromosomes are no longer masked. Relative to the model with dosage compensation, we have weaker allelic and epistatic effects of the X chromosome. Hence, incompatible island X alleles are easier go get swamped and also have a more difficult time to keep incompatible continental (A or X) alleles from swamping.

The consequences can most easily be seen in the first row of Figure 3(a)-(c) with pure female migration, where, in contrast to dosage compensation, differences between the various genomic architectures are not compensated anymore. We observe a strong asymmetry between 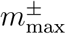 of X→A and A→X-models for all levels of male migration relative to the corresponding results with dosage compensation. Here migration bounds for X→A always exceed those obtained for A→X-models. Intuitively, one can understand this as follows: In the X→A model, three immigrating X chromosomes “fight” against four resident autosomes, whereas in A→X the odds are in favor of the immigrating autosomes. Thus the island is swamped more easily in the latter case.

As seen for dosage compensation before, for weak epistasis (γ ≈ 0, pure “selection against immigrants”), it is always the locus with weaker direct effect that is swamped first. In our example this is always the “continental” locus, because we have stronger selection on the island locus. For unbiased migration (Fig. 3(f)) all models converge to the same bound. However, introducing sex-biased migration leads to relative higher gene flow on the X for a female bias (and therefore lower bounds for models with immigrating X), as can be seen in Figure 3(c). Similarly, male-biased migration leads to weaker X-linked gene flow and a higher bounds in these models, *i.e.* X→A and X→X, (Fig. 3(i)).

If we compare migration bounds of Figure 2 and 3, we can see that dosage compensation outbalances most of the differences in 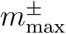 between A→X and X→A, especially for local stability. While dosage compensation strengthens the fitness effect of the island allele in A→X, the increased epistatic pressure on the continental allele in X→A is outbalanced by its increased fitness effect.

### 3.2 Cytonuclear (mitochondrial) codominant DMIs

Finally, we investigate cytonuclear DMIs in Figure 4, where a gene in the haploid mitochondrial genome (termed o/O for organelle) is incompatible with a nuclear locus. Dosage compensation of the X chromosome again means that the male XyO-hybrids suffer as much as the female XXO-hybrids while they suffer only as much as XxO hybrids without dosage compensation. Relative to nuclear DMIs, three main effects lead to changes in 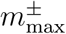:

First, the cytoplasmic locus experiences effectively stronger direct and epistatic selection (factor two in Eqs. (B.34c)), because we maintain the per locus effect identical to nuclear loci. Since a single allele already accounts for the full mitochondrial locus effect this leads to a larger effect per allele. As a consequence, 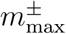 without sex-bias in migration is elevated relative to A→A model (gray lines in Fig. 4(a)-(c)).

Second, sex-biased migration has an even stronger effect in cytonuclear DMIs than in the X-linked nuclear DMIs: Since mitochondria are maternally inherited, the effective gene flow for mitochondrial loci is reduced to zero with pure male migration. Consequently, all migration bounds with immigrating incompatible mitochondrial genes diverge to infinity. In Figure 4 (last two rows), we study the case of strong, but not complete male bias (*R* = *−*0.9). Since the migration pressure on the mitochondrial locus and the X chromosome is reduced, migration bounds 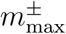 increase for all cytonuclear DMIs, especially for those also involving X chromosomes. This increase in 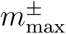 is even further promoted by dosage compensation, strengthening the effect of X.

Finally, because of strict maternal inheritance, the dynamics of the mitochondrial locus is not influenced by any fitness effects in males. In *𝒳*-*𝒪* models this also entails that dosage compensation only affects the dynamics of the X-locus - in contrast to nuclear DMIs, where also autosomal loci are affected if they interact with a hemizygous X locus. As a consequence, the boosting effect of dosage compensation on 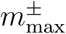 is symmetric for O→X and X→O, in stark contrast to nuclear DMIs, where dosage compensation does not change much for X→A while it strongly increases A→X.

**Figure 4:**
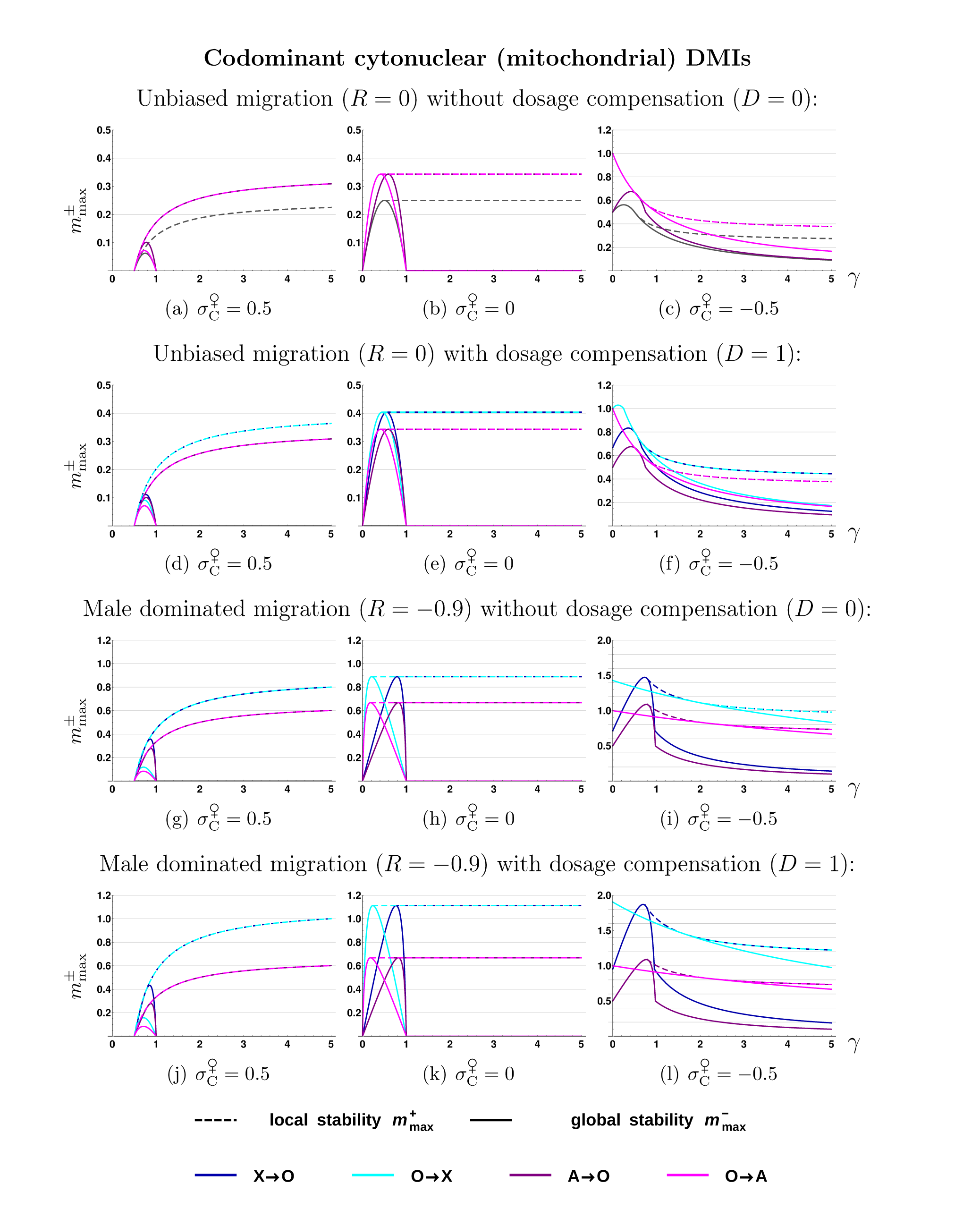
Codominant cytonuclear DMIs. Maximum permissible migration rates for local stability either coincide for all models (a)-(c), or just for X→O and O→X as well as for O→A and A→O in all other cases. Migration bounds for global stability only coincide without dosage compensation or sex-biased migration between O→X and O→A, as well as for X→O and A→O. The A→A model is given in panel (a)-(c) in gray as a reference. All bounds 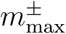 are derived analytically, see Eqs.(B.40),(B.42). See Figure 2 for further explanations. Note the different scaling of the y-axis in third column.

## 4 Discussion

If barriers to gene flow build up among populations in primary or secondary contact, this can have important consequences for their genetic architecture. A lot of recent interest has focused on *islands of speciation* (*or divergence*) (Wu, 2001; Turner et al., 2005; Butlin et al., 2012; Nosil, 2012; Nosil and Feder, 2012; Via, 2012), yet corresponding empirical findings are equivocal on that matter (Cruickshank and Hahn, 2014; Pennisi, 2014). There are, however, several clear and undisputed genomic patterns of speciation, on which we concentrate here. The most widely known ones are Haldane’s rule, (Haldane, 1922), which has motivated much previous speciation research (see reviews and examples in Coyne and Orr (2004); Presgraves (2008); Lachance and True (2010); Presgraves (2010); Oka and Shiroishi (2013) and the *large X*-*effect* (reviewed in Presgraves, 2008), which both highlight an important role of the X chromosome (or the Z chromosome in birds) in speciation. In addition, hybrid incompatibilities are frequently observed also between nuclear and cytoplasmic markers. Plants show incompatibilities with plastid genomes (Greiner et al., 2011; Snijder et al., 2007) and mitochondria have been reported to be incompatible with nuclear genes across a wide range of species (Ellison and Burton, 2008; Lee et al., 2008; Burton and Barreto, 2012). In insects, cytoplasmic incompatibilities can also be caused by infections with the intracellular bacterium Wolbachia (O’Neill et al., 1992; Werren, 1997; Coyne and Orr, 2004).

In the current study we investigate how the genetic architecture of an inital hybrid incompatibility between incipient sister species can maintain divergence in the presence of ongoing gene flow. Can (primary or secondary) gene flow favor X-linked or cytonuclear DMIs over autosomal ones, and if so under which conditions? We studied this question about a possible first step towards speciation using a minimal model of a two-locus DMI in a continent-island population that allows for analytical treatment. We derive maxium permissible migration bounds which still permit maintenance of a DMI in the face of gene flow. Conditions that yield increased migration limits facilitate speciation, as they are lost less easily and can subsequently provide more persistent seeds for further ongoing differentiation.

### Conditions for parapatric DMIs

Like in the autosomal case (Bank et al., 2012), the origin and maintenance of a two-locus X-linked or cytonuclear DMI requires that at least one of the DMI substitutions (namely: the incompatible variant on the island) is adaptive. If multi-locus barriers to gene flow build up gradually from initial two-locus incompatibilities, this confirms that postzygotic parapatric speciation requires at least some degree of ecological differentiation and local adaptation. Empirically, there is widespread evidence for positive selection on genes involved in DMIs (Macnair and Christie, 1983; Ting et al., 1998; Presgraves et al., 2003; Barbash et al., 2004; Dettman et al., 2007).

For all types of DMIs, we observe two basic selective forces driving their evolution. Selection against immigrants implies that the new migrants have a fitness deficit relative to island residents, resulting in *ecological speciation* scenarios (Schluter and Conte, 2009; Nosil, 2012). A characteristic of this regime is that evolution of a stable DMI is independent of its evolutionary history.

Alternatively, a stable DMI is caused by selection against hybrids, where migrants can even have a positive fitness. If hybrids are unfit, immigrants still suffer an indirect disadvantage as long as they are rare and their genotypes are readily broken down by sex and recombination. This scenario typically leads to a bistable dynamics, where a stable DMI will only evolve from favorable starting conditions or permissive evolutionary histories (such as secondary contact). The scenario has also been referred to as *mutation-order-speciation* (Mani and Clarke, 1990).

We measure the strength of a parapatric DMI by means of two migration bounds. The higher one, 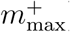, is the limit beyond which a DMI can neither evolve nor an existing one can be maintained. The lower bound, 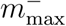, is the limit up to which a DMI will always evolve in the face of gene flow, irrespective of the evolutionary history (globally stable DMI). For migration rates between both bounds, a DMI is maintained, but will evolve only under favorable histories, such as secondary contact, or if the second incompatible substitution occurs on the continent.

### Contrasting different DMI architectures

We find that the genetic architecture of a DMI (with incompatible genes on autosomes, X chromosomes, or in the mitochondrial genome) can have a strong effect on its stability. However, this effect also crucially depends on other factors, such as, in particular, the level of dosage compensation and the sex-bias in the migration rates.

First, without dosage compensation and without sex-biased gene flow, the hemizygosity of the X chromosome in males leads to shifts of 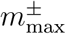 in the presence of epistasis compared to autosome-autosome DMIs. This is due to ploidy differences: “3 X chromosomes fight 4 autosomes”. Therefore, the A→X scenario (where a resident X-linked allele competes with an immigrating incompatible autosomal gene) constitutes a weaker barrier to gene flow than the X→A model. Note that this effect depends crucially on the (negative) epistasis of the DMI and is not observed in a single-locus model of local adaptation. Second, dosage compensation strengthens the X alleles, which leads to higher migration bounds in all X-linked DMIs. In particular, it increases stability of DMIs with an incompatible X locus on the island, compensating the A→X versus X→A asymmetry. Third, sex-biased migration leads to lower/higher limits for DMIs with immigrating X for female/male bias. Fourth, our results in the SI Section A.1 show no large difference between codominant and recessive nuclear DMIs (which lead to Haldane’s rule) concerning the migration bounds. In fact, the difference for X-linked DMIs are even smaller than for autosome-autosome DMIs. Fifth, for cytonuclear DMIs we often observe stronger barriers to gene flow since the haploid cytoplasmic alleles experience the full locus effect. Furthermore, sex-bias in migration yields an especially strong effect, as for pure male migration effective gene flow at the mitochondrial locus ceases completely.

Our numerical simulations for the effect of LD in the SI Section A.2 agree with the approximate analytical results for weak and moderately strong DMIs. For very strong DMIs, stronger deviations occur for codominant A→A and X→X DMIs, which maintain very strong LD once all (male and female) hybrids with incompatible alleles are almost inviable/infertile. As a consequence, gene flow among the continent and island haplotypes is blocked and we obtain higher migration bounds relative to X→A and A→X DMIs. For the latter two, F1 hybrid males carrying the compatible x allele (genotype Aaxy) are viable and can produce ax gametes for the F2 generation. This effect of extreme LD and blocked gene flow does not exist for recessive DMIs (see SI Section A.1 for details). Our numerical simulations also show that the effect of drift is usually small and does not lead to qualitative changes (SI Section A.3). Since DMI alleles can be lost by drift, stochastic migration bounds 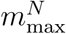 are generally smaller than their deterministic counterparts. In SI Section A.3, we present an analytical approximation to estimate this reduction due to drift.

### The *large X*-*effect*

Summarizing all different cases described above, we find that the most stable DMIs are almost always X-linked, where migration bounds are typically enhanced by a factor of 4/3 to 2 relative to autosomal DMIs (unless migration is strongly female biased). Although this is not a very strong effect, it is very general and applies whenever gene flow plays a role at any stage of the speciation process. This includes, in particular, scenarios of secondary contact and also later stages of the speciation process where additional barriers to gene flow exist in the genomic background. In this latter case, the gene flow at the focal DMI loci needs to be replaced by appropriate effective migration rates (Barton and Bengtsson, 1986). The pattern that follows from a more stable X barrier is consistent with a higher density of X-linked hybrid incompatibilities, the *large X*-*effect*.

Our results show a clear boost of X migration bounds, in particular, if there is dosage compensation and if migration is male biased. Empirical studies show that sex-biased migration is common in nature and report a prevalence for migration of the heterogametic sex in both mammals, where dispersal is on average male biased, (Lawson Handley and Perrin, 2007) and in birds, where female dispersal dominates (Greenwood, 1980). In the context of our results, these trends strengthen the predicted pattern of a *large X*-*effect* or *large Z*-*effect*, respectively.

One example stems from the house mouse, *Mus musculus*. There is strong empirical evidence for a *large X*-*effect* in this species (Tucker et al., 1992; Good et al., 2008; White et al., 2012), such as the major involvement of the X chromosome in hybrid sterility (Oka et al., 2004; Storchova et al., 2004). Mice exhibit rather complete dosage compensation due to X-inactivation in females (Payer and Lee, 2008). Furthermore, the house mouse displays male-biased dispersal at breeding age (Greenwood, 1980; Gerlach, 1990).

Several alternative mechanisms as potential underlying causes for a *large X*-*effect* have been discussed in the literature, such as sex ratio meiotic drive, regulation of the X chromosome in the male germ line (Coyne and Orr, 2004; Presgraves, 2008), or faster evolution of the X chromosome (termed *faster-X-effect* Charlesworth et al., 1987). In the panmictic population model by Charlesworth et al. (1987), faster evolution on the X chromosome results if adaptations are, on average, recessive and are thus exposed to stronger selection on the hemizygous X. We note that our model with gene flow predicts an advantage of X-linked genes for island adaptations even if they are not recessive, but codominant (or even slightly dominant, see SI Section A.4 for details and proofs). Since the *faster X-effect* (more adaptations on the X) also favors a *larger X-effect* (more incompatibilities involving the X), this is another way how speciation with gene-flow can contribute to this pattern. In summary a mono-causal explanation for the *large X*-*effect* seems unlikely, and it remains an open question, to which extent each factor contributes. Our study adds differentiation under gene flow as another element to this mix.

Our results relate to Haldane’s rule only in so far as this pattern partially overlaps with the *large X*-*effect*. Beyond that, we do not obtain a prediction. In particular, the migration bounds for codominant and recessive DMIs are similar (while only the latter lead to Haldane’s rule).

### Introgression patterns

A second conclusion from our results that can be related to data is that X-linked alleles in an incompatibility face stronger barriers to introgression than the corresponding autosomal alleles. This effect rests on two basic observations: the tendency for higher migration bounds of all X-linked DMIs with dosage compensation (which also contributes to a large X-effect), and the asymmetry promoting A→X over X→A introgression that we observe for the incompatible allele if dosage compensation is incomplete or absent (the 3 versus 4 chromosomes effect). Our findings agree with the result of a recent simulation study for DMIs on a cline by Wang (2013), who showed that, for an X-autosome DMIs, the incompatible X allele flows less easily across a cline than the autosomal allele.

A pattern of reduced X-introgression relative to autosomal introgression has been recognized in many sister-species in nature. In the complex of *Anopheles gambiae* sister clades Fontaine et al. (2015) found “pervasive autosomal introgression” between different species, in contrast to the X chromosome, which contains disproportionately more factors in reproductive isolation.

Liu et al. (2015) report three interspecies hybridization events in mice (*Mus musculus domesticus* and *M. spretus*), leading to exclusively autosomal, partially adaptive introgression. Similarly, Macholán et al. (2007) showed weaker introgression patterns and lower selection pressure on the X chromosomes compared to the autosomes in the central European mouse hybrid zone of *Mus musculus musculus* and *M. m. domesticus*. The authors suppose that the X is shielded more effectively from introgression due to the *large X*-*effect*.

Further examples exist for birds. Sætre et al. (2003) report “rather extensive hybridization and backcrossing in sympatry” between two populations of flycatchers hybridizing in secondary contact. Nevertheless, gene flow was again predominantly found on the autosome. Hooper and Price (2015) report that derived cross-species inversions among sister species of Estrilid finches are strongly enriched on the Z chromosome. The pattern is strongest in continental clades with high level of sympatry and (plausibly) higher levels of gene-flow during the speciation process. If inversions harbor DMIs, this is consistent with our finding that derived incompatibilities on the Z chromosome are more stable to gene flow than autosomal incompatibilities.

Also other factors, such as recombination, can influence differential introgression on X chromosomes and autosomes. Indeed, there is empirical evidence that recombination can structure autocorrelation patterns among introgressed loci. However, available data also show that recombination cannot be the the sole explanation for differential introgression among genomic regions, e.g. in mice (Payseur et al., 2004) or finches (Hooper and Price, 2015). As for the *large X*-*effect* our mechanism is one of several possible ones.

### Biological assumptions and limitations of the model

Our study has been intended as a minimal model approach that allows for analytical treatment. As such, it rests on several simplifying assumptions concerning the genetics of the DMI and the ecological setting. These limitations suggest possible model extensions for future work.

All our results assume a simple DMI between just two loci. This is in line with most previous theoretical work and known empirical cases (Coyne and Orr, 2004; Maheshwari and Barbash, 2011). Nevertheless, complex DMIs involving multiple loci are clearly relevant at later stages of a speciation process and could lead to new effects that are not captured here (*e.g.* Lindtke and Buerkle, 2015).

Our fitness scheme for two-locus DMIs comprises codominant and recessive cases. Empirically, the functional form depends on the underlying mechanisms causing hybrid fitness loss, which is still debated. Hybrid incompatibilities can be due to loss-of-function or gain-of-function mutations (reviewed by Maheshwari and Barbash, 2011). While the former tend to act recessively, the latter will likely affect heterozygotes, and may be better captured by a partially dominant DMI.

Recessive DMIs, in turn, occur in a number of different types, (*e.g.* Presgraves, 2010; Cattani and Presgraves, 2012; Matsubara et al., 2015), which lead to slightly different models. We have briefly studied some of these alternatives analytically, such as a *recessive-A codominant-X* -DMI or a *codominant-A recessive-X* -DMI (data not shown). We did not detect any noteworthy difference in their evolutionary dynamics or for the migration bounds relative to the results reported here. Still, more relevant changes are clearly possible, for example if the single locus effects can lead to over-or underdominance.

For the results presented, we assume that dosage compensation enhances not only the single-locus effect, but also the incompatibility. Empirically, hybrid incompatibilities are frequently dosage-sensitive, *e.g.* in a *Arabidopsis thaliana*/ *A. arenosa* cross, where a DMI results due to failure in gene silencing Josefsson et al. (2006), or in a *Mus musculus musculus*/ *M. m. domesticus* cross, where X-linked hybrid male sterility results from over-expression of X-linked genes in spermatogenesis (Good et al., 2010). Furthermore, in haplo-diploid Nasonian wasps genetically engineered diploid males were less affected by hybrid sterility than haploid male hybrids, pointing to a strong effect of ploidy on hybrid fertility Beukeboom et al. (2015).

Nevertheless, we also investigated the effect of dosage compensation only on the single locus effect or only on the incompatibility (results not shown). As expected, we obtain intermediate patterns between no and full dosage compensation.

Concerning the ecological assumptions, our model assumes unidirectional gene flow between two panmictic demes. While our results readily extend to weak back migration (which leads only to slight shifts of the equilibria), strong bidirectional migration can lead to qualitatively new effects that are not captured by our framework. For example, polymorphisms at single loci can be maintained for arbitrarily strong gene flow if heterogeneous soft selection leads to a rare-type advantage (Levene, 1953). Furthermore, generalist genotypes that are inferior in both demes, but do well on average, can be maintained if (and only if) bidirectional migration is sufficiently strong (see Akerman and Bürger, 2014, for results in a two-locus model without epistasis).

Alternative models for the population structure can also lead to substantial differences. In particular, our two-deme model ignores isolation by distance, which can be captured either in a discrete cline model with a chain of demes, or in a continuous-space framework. It is expected that polymorphisms (and DMIs) can be maintained with much larger gene flow (or weaker selection) in these settings (Barton, 2013). Still, several of our key results, such as reduced introgression of X-linked incompatibility alleles, should still hold under these conditions (see Wang, 2013, for a discrete cline model).

## 5 Acknowledgements

We thank Andrea Betancourt, Alexandre Blanckaert, Reinhard Bürger, Brian Charlesworth, Andy Clark, Sebastian Matuszewsky, Sylvain Mousset, Mohamed Noor, Sally Otto, Christian Schlötterer, Maria Servedio, Derek Setter, Claus Vogl and three referees for helpful discussions, suggestions and comments on the manuscript. This work was made possible with financial support by the Austrian Science Fund (FWF) via funding for the Vienna Graduate School for Population Genetics.

## References

Agrawal, A., Feder, J., and Nosil, P. (2011). Ecological divergence and the origins of intrinsic postmating isolation with gene flow. International Journal of Ecology, 2011.

Akerman, A. and Bürger, R. (2014). The consequences of gene flow for local adaptation and differentiation: a two-locus two-deme model. Journal of mathematical biology, 68(5):1135–1198.

Bank, C., Bürger, R., and Hermisson, J. (2012). The limits to parapatric speciation: Dobzhansky–Muller incompatibilities in a continent–island model. Genetics, 191(3):845–863.

Barbash, D., Awadalla, P., and Tarone, A. (2004). Functional divergence caused by ancient positive selection of a Drosophila hybrid incompatibility locus. PLoS biology, 2:839–848.

Barnard-Kubow, K. B., So, N., and Galloway, L. F. (2016). Cytonuclear incompatibility contributes to the early stages of speciation. Evolution, 70(12):2752–2766.

Barton, N. (2013). Does hybridization influence speciation? Journal of evolutionary biology, 26(2):267–269.

Barton, N. and Bengtsson, B. (1986). The barrier to genetic exchange between hybridising populations. Heredity, 57(3):357–376.

Bateson, W. (1909). Heredity and variation in modern lights. Darwin and modern science, 85:101.

Beukeboom, L., Koevoets, T., Morales, H. E., Ferber, S., and van de Zande, L. (2015). Hybrid incompatibilities are affected by dominance and dosage in the haplodiploid wasp Nasonia. Frontiers in Genetics, 6:14.

Bürger, R. and Akerman, A. (2011). The effects of linkage and gene flow on local adaptation: A two-locus continent–island model. Theoretical population biology, 80(4):272–288.

Burton, R. and Barreto, F. (2012). A disproportionate role for mtDNA in Dobzhansky–Muller incompatibilities? Molecular Ecology, 21(20):4942–4957.

Butlin, R., Debelle, A., Kerth, C., Snook, R., Beukeboom, L., Castillo, C., Diao, W., Maan, M., Paolucci, S., Weissing, F., et al. (2012). What do we need to know about speciation? Trends in Ecology & Evolution, 27(1):27–39.

Cattani, M. and Presgraves, D. (2012). Incompatibility Between X Chromosome Factor and Pericentric Heterochromatic Region Causes Lethality in Hybrids Between Drosophila melanogaster and Its Sibling Species. Genetics, 191(2):549–559.

Charlesworth, B., Coyne, J., and Barton, N. (1987). The relative rates of evolution of sex chromosomes and autosomes. American Naturalist, pages 113–146.

Corbett-Detig, R., Zhou, J., Clark, A., Hartl, D., and Ayroles, J. (2013). Genetic incompatibilities are widespread within species. Nature, 504(7478):135–137.

Coyne, J. and Orr, H. (1989). Two rules of speciation. Speciation and its Consequences, pages 180–207.

Coyne, J. and Orr, H. (2004). Speciation. Sinauer Associates Sunderland, MA.

Cruickshank, T. and Hahn, M. (2014). Reanalysis suggests that genomic islands of speciation are due to reduced diversity, not reduced gene flow. Molecular Ecology, 23(13):3133–3157.

Dettman, J., Sirjusingh, C., Kohn, L., and Anderson, J. (2007). Incipient speciation by divergent adaptation and antagonistic epistasis in yeast. Nature, 447(7144):585–588.

Dobzhansky, T. (1936). Studies on hybrid sterility. II. Localization of sterility factors in Drosophila pseudoobscura hybrids. Genetics, 21(2):113.

Ellegren, H. (2009). Genomic evidence for a large-Z effect. Proceedings of the Royal Society B: Biological Sciences, 276(1655):361–366.

Ellegren, H., Hultin-Rosenberg, L., Brunström, B., Dencker, L., Kultima, K., and Scholz, B. (2007). Faced with inequality: chicken do not have a general dosage compensation of sex-linked genes. BMC biology, 5(1):40.

Ellison, C. and Burton, R. (2008). Interpopulation hybrid breakdown maps to the mitochondrial genome. Evolution, 62(3):631–638.

Feder, J. and Nosil, P. (2009). Chromosomal inversions and species differences: when are genes affecting adaptive divergence and reproductive isolation expected to reside within inversions? Evolution, 63(12):3061–3075.

Felsenstein, J. (1981). Skepticism towards Santa Rosalia, or why are there so few kinds of animals? Evolution, pages 124–138.

Fontaine, M., Pease, J., Steele, A., Waterhouse, R. M., Neafsey, D., Sharakhov, I., Jiang, X., Hall, A., Catteruccia, F., Kakani, E., et al. (2015). Extensive introgression in a malaria vector species complex revealed by phylogenomics. Science, 347(6217):1258524.

Gavrilets, S. (1997). Hybrid zones with Dobzhansky-type epistatic selection. Evolution, pages 1027–1035.

Gerlach, G. (1990). Dispersal mechanisms in a captive wild house mouse population (Mus domesticus Rutty). Biological Journal of the Linnean Society, 41(1-3):271–277.

Good, J., Dean, M., and Nachman, M. (2008). A complex genetic basis to X-linked hybrid male sterility between two species of house mice. Genetics, 179(4):2213–2228.

Good, J., Giger, T., Dean, M., and Nachman, M. (2010). Widespread over-expression of the X chromosome in sterile F1 hybrid mice. PLoS genetics, 6(9):e1001148.

Graves, J., Disteche, C., et al. (2007). Does gene dosage really matter. J Biol, 6(1):1.

Greenwood, P. (1980). Mating systems, philopatry and dispersal in birds and mammals. Animal behaviour, 28(4):1140–1162.

Greiner, S., Rauwolf, U., Meurer, J., and Herrmann, R. (2011). The role of plastids in plant speciation. Molecular ecology, 20(4):671–691.

Haldane, J. (1922). Sex ratio and unisexual sterility in hybrid animals. Journal of genetics, 12(2):101–109.

Hooper, D. and Price, T. (2015). Rates of karyotypic evolution in Estrildid finches differ between island and continental clades. bioRxiv, page 013987.

Josefsson, C., Dilkes, B., and Comai, L. (2006). Parent-dependent loss of gene silencing during interspecies hybridization. Current Biology, 16(13):1322–1328.

Kimura, M. (1957). Some problems of stochastic processes in genetics. The Annals of Mathematical Statistics, pages 882–901.

Lachance, J. and True, J. (2010). X-autosome incompatibilities in Drosophila melanogaster: tests of Haldane’s rule and geographic patterns within species. Evolution, 64(10):3035–3046.

Lawson Handley, L. and Perrin, N. (2007). Advances in our understanding of mammalian sex-biased dispersal. Molecular Ecology, 16(8):1559–1578.

Lee, H., Chou, J., Cheong, L., Chang, N., Yang, S., and Leu, J. (2008). Incompatibility of nuclear and mitochondrial genomes causes hybrid sterility between two yeast species. Cell, 135(6):1065–1073.

Levene, H. (1953). Genetic equilibrium when more than one ecological niche is available. American Naturalist, pages 331–333.

Lindtke, D. and Buerkle, C. (2015). The genetic architecture of hybrid incompatibilities and their effect on barriers to introgression in secondary contact. Evolution.

Liu, K., Steinberg, E., Yozzo, A., Song, Y., Kohn, M., and Nakhleh, L. (2015). Interspecific introgressive origin of genomic diversity in the house mouse. Proceedings of the National Academy of Sciences, 112(1):196–201.

Macholán, M., Munclinger, P., Šugerková, M., Dufková, P., Bímová, B., Božíková, E., Zima, J., and Piálek, J. (2007). Genetic analysis of autosomal and X-linked markers across a mouse hybrid zone. Evolution, 61(4):746–771.

Macnair, M. and Christie, P. (1983). Reproductive isolation as a pleiotropic effect of copper tolerance in Mimulus guttatus. Heredity, 50(3):295–302.

Maheshwari, S. and Barbash, D. (2011). The genetics of hybrid incompatibilities. Annual review of genetics, 45:331–355.

Mallet, J. (2008). Hybridization, ecological races and the nature of species: empirical evidence for the ease of speciation. Philosophical Transactions of the Royal Society B: Biological Sciences, 363(1506):2971–2986.

Mani, G. and Clarke, B. (1990). Mutational order: a major stochastic process in evolution. Proceedings of the Royal Society of London B: Biological Sciences, 240(1297):29–37.

Masly, J. and Presgraves, D. (2007). High-resolution genome-wide dissection of the two rules of speciation in Drosophila. PLoS Biol, 5(9):e243.

Matsubara, K., Yamamoto, E., Mizobuchi, R., Yonemaru, J., Yamamoto, T., Kato, H., and Yano, M. (2015). Hybrid Breakdown Caused by Epistasis-Based Recessive Incompatibility in a Cross of Rice (Oryza sativa L.). Journal of Heredity, 106(1):113–122.

Muller, H. (1942). Isolating mechanisms, evolution and temperature. In Biological Symposia, volume 6, pages 71–125.

Nosil, P. (2012). Ecological speciation. Oxford University Press.

Nosil, P. and Feder, J. (2012). Genomic divergence during speciation: causes and consequences. Philosophical Transactions of the Royal Society B: Biological Sciences, 367(1587):332–342.

Oka, A., Mita, A., Sakurai-Yamatani, N., Yamamoto, H., Takagi, N., Takano-Shimizu, T., Toshimori, K., Moriwaki, K., and Shiroishi, T. (2004). Hybrid breakdown caused by substitution of the X chromosome between two mouse subspecies. Genetics, 166(2):913–924.

Oka, A. and Shiroishi, T. (2013). Regulatory divergence of X-linked genes and hybrid male sterility in mice. Genes & genetic systems, 89(3):99–108.

O’Neill, S., Giordano, R., Colbert, A., Karr, T., and Robertson, H. (1992). 16S rRNA phylogenetic analysis of the bacterial endosymbionts associated with cytoplasmic incompatibility in insects. Proceedings of the National Academy of Sciences, 89(7):2699–2702.

Orr, H. and Turelli, M. (2001). The evolution of postzygotic isolation: accumulating Dobzhansky-Muller incompatibilities. Evolution, 55(6):1085–1094.

Payer, B. and Lee, J. (2008). X chromosome dosage compensation: how mammals keep the balance. Annual review of genetics, 42:733–772.

Payseur, B. A., Krenz, J. G., and Nachman, M. W. (2004). Differential patterns of introgression across the x chromosome in a hybrid zone between two species of house mice. Evolution, 58(9):2064–2078.

Pennisi, E. (2014). Disputed islands. Science, 345(6197):611–613.

Presgraves, D. (2008). Sex chromosomes and speciation in Drosophila. Trends in Genetics, 24(7):336–343.

Presgraves, D. (2010). The molecular evolutionary basis of species formation. Nature Reviews Genetics, 11(3):175–180.

Presgraves, D., Balagopalan, L., Abmayr, S., and Orr, H. (2003). Adaptive evolution drives divergence of a hybrid inviability gene between two species of Drosophila. Nature, 423(6941):715–719.

Rutschman, D. H. (1994). Dynamics of the two-locus haploid model. Theoretical population biology, 45(2):167–176.

Sætre, G., Borge, T., Lindroos, K., Haavie, J., Sheldon, B., Primmer, C., and Syvänen, A.-C. (2003). Sex chromosome evolution and speciation in Ficedula flycatchers. Proceedings of the Royal Society of London B: Biological Sciences, 270(1510):53–59.

Schluter, D. and Conte, G. (2009). Genetics and ecological speciation. Proceedings of the National Academy of Sciences, 106(Supplement 1):9955–9962.

Slatkin, M. (1987). Gene flow and the geographic structure of natural populations. Science, 236:787–792.

Snijder, R., Brown, F., and van Tuyl, J. (2007). The role of plastome-genome incompatibility and biparental plastid inheritance in interspecific hybridization in the genus Zantedeschia (Araceae). Floriculture and Ornamental Biotechnology, 1(2):150–157.

Storchova, R., Gregorová, S., Buckiova, D., Kyselova, V., Divina, P., and Forejt, J. (2004). Genetic analysis of X-linked hybrid sterility in the house mouse. Mammalian Genome, 15(7):515–524.

Sweigart, A. and Flagel, L. (2014). Evidence of Natural Selection Acting on a Polymorphic Hybrid Incompatibility Locus in Mimulus. Genetics, pages genetics–114.

Ting, C., Tsaur, S., Wu, M., and Wu, C. (1998). A rapidly evolving homeobox at the site of a hybrid sterility gene. Science, 282(5393):1501–1504.

Tucker, P., Sage, R., Warner, J., Wilson, A., and Eicher, E. (1992). Abrupt cline for sex chromosomes in a hybrid zone between two species of mice. Evolution, pages 1146–1163.

Turelli, M. and Orr, H. (2000). Dominance, epistasis and the genetics of postzygotic isolation. Genetics, 154(4):1663–1679.

Turner, T., Hahn, M., and S., N. (2005). Genomic islands of speciation in Anopheles gambiae. PLoS biology, 3(9):e285.

Via, S. (2012). Divergence hitchhiking and the spread of genomic isolation during ecological speciation-with-gene-flow. Philosophical Transactions of the Royal Society B: Biological Sciences, 367(1587):451–460.

Via, S. and West, J. (2008). The genetic mosaic suggests a new role for hitchhiking in ecological speciation. Molecular Ecology, 17(19):4334–4345.

Wang, R. (2013). Gene flow across a hybrid zone maintained by a weak heterogametic incompatibility and positive selection of incompatible alleles. Journal of Evolutionary Biology, pages 386–396.

Werren, J. (1997). Biology of wolbachia. Annual review of entomology, 42(1):587–609.

White, M., Stubbings, M., Dumont, B. L., and Payseur, B. (2012). Genetics and evolution of hybrid male sterility in house mice. Genetics, 191(3):917–934.

Wu, C. (2001). The genic view of the process of speciation. Journal of Evolutionary Biology, 14(6):851–865.

Yeaman, S. and Otto, S. P. (2011). Establishment and maintenance of adaptive genetic divergence under migration, selection, and drift. Evolution, 65(7):2123–2129.

